# Redox regulation by TXNRD3 during epididymal maturation underlies capacitation-associated mitochondrial activation and sperm motility in mice

**DOI:** 10.1101/2021.07.19.452987

**Authors:** Huafeng Wang, Qianhui Dou, Kyung Jo Jung, Jungmin Choi, Vadim N. Gladyshev, Jean-Ju Chung

**Author notes:** **Correspondence:** Jean-Ju Chung.

## Abstract

During epididymal transit, redox remodeling protects mammalian spermatozoa, preparing them for survival in the subsequent journey to fertilization. However, molecular mechanisms of redox regulation in sperm development and maturation remain largely elusive. In this study, we report that TXNRD3, a thioredoxin reductase family member particularly abundant in elongating spermatids at the site of mitochondrial sheath formation, regulates redox homeostasis to support male reproduction. Using *Txnrd3*^*-/-*^ mice, our biochemical, ultrastructural, and live cell imaging analyses revealed impairments in sperm morphology and motility under conditions of TXNRD3 deficiency. We reveal that mitochondria develop more defined cristae during capacitation in wild type sperm. Absence of TXNRD3 alters redox status in both the head and tail during sperm maturation and capacitation, resulting in defective mitochondrial ultrastructure and activity under capacitating conditions. These findings provide insights into molecular mechanisms of redox homeostasis and bioenergetics during sperm maturation, capacitation, and fertilization.

## Introduction

Testicular sperm are functionally immature in that they lack the ability to naturally fertilize an egg. Epididymis transit is an indispensable step for mammalian sperm cells to fully develop the fertilizing ability. During epididymal descent, spermatozoa acquire the potential to develop progressive motility and capacitation (Sullivan and Mieusset 2016, Gervasi and Visconti 2017). One major threat to sperm for the subsequent journey to fertilization is oxidative damage since sperm plasma membrane is rich in polyunsaturated fatty acids (Vernet et al. 2004). Redox mechanisms prepare epididymal spermatozoa for their protection and survival during the fertilization journey (Chabory et al. 2009). For example, oxidation of the intracellular milieu is involved in mammalian sperm maturation, capacitation, acrosome reaction (de Lamirande et al. 1997, O’Flaherty 2015). At the same time, oxidative damage correlates with lower motility of human sperm (Kao et al. 2008) and causes DNA fragmentation and protein crosslinking (Henkel et al. 2005). However, it remains unclear how redox homeostasis is maintained during mammalian sperm capacitation.

Thioredoxin and glutathione systems are two major redox systems that utilize the thiol redox biology to maintain cellular redox homeostasis and protect against oxidative stress. Thioredoxin reductases, a family of selenium-containing pyridine nucleotide-disulfide oxidoreductases, are key redox regulators of mammalian thioredoxin system. They are comprised of three enzymes: thioredoxin reductase 1 (TXNRD1), thioredoxin reductase 2 (TXNRD 2) (Holmgren 1980, Mustacich and Powis 2000) and thioredoxin-glutathione reductase (TXNRD3, also known as TGR) (Sun et al. 1999, Sun et al. 2001). TXNRD1 and TXNRD2 are essential proteins that support redox homeostasis in the cytosol and mitochondria, respectively (Arnér and Holmgren 2000, Conrad et al. 2004, Nalvarte et al. 2004, Jakupoglu et al. 2005). TXNRD3 is predominantly expressed in testis (Sun et al. 2001) and particularly abundant in elongating spermatids at the site of mitochondrial sheath formation (Su et al. 2005). Structurally, TXNRD3 contains an additional N-terminal glutaredoxin domain compared with the other two TXNRD isozymes (Sun et al. 2001). Disulfide bond isomerase activity of TXNRD3 was previously reported *in vitro* (Su et al. 2005), yet the physiological significance and underlying mechanisms of TXNRD3 in male germ cell development and sperm function have been largely unknown. In the accompanying study, we report that *Txnrd3*^*-/-*^ mice exhibit compromised male fertility, with mild impregnation delay *in vivo* and diminished sperm fertilization capacity *in vitro* (Dou et al., 2021).

Here, we report the underlying mechanism of TXNRD3 function in regulation of chromatin organization and mitochondrial activity during sperm capacitation in mice. We find that targeted disruption of *Txnrd3* induces capacitation-associated impairment in sperm morphology and motility. Biochemical analyses demonstrate that TXNRD3 expression diminishes during sperm maturation, impairing redox homeostasis of head and flagellar proteins. Using ultrastructural analyses and live cell imaging, we reveal that mitochondria in *Txnrd3*^*-/-*^ sperm undergo structural defects and lose control of its activity especially during sperm capacitation. These findings provide insights into sperm redox regulation during epididymal maturation and its effect on capacitation-associated energy metabolism.

## Materials and methods

### Mouse Sperm Preparation and *in vitro* Capacitation

Epididymal spermatozoa from adult male mice were collected by swim-out from caudal epididymis in standard HEPES saline (HS) medium (in mM, 135 NaCl, 5 KCl, 1 MgSO_4_, 2 CaCl_2_, 20 HEPES, 5 Glucose, 10 Lactic acid, 1 Na pyruvate, pH 7.4 adjusted with NaOH, 320 mOsm/L). 2×10^6^ cells/ml sperm were incubated in human tubular fluid (HTF) medium (EMD Millipore) at 37°C, 5% CO_2_ for 90 min to induce capacitation.

### Antibodies and Reagents

Rabbit polyclonal antibodies specific to mouse TXNRD3 and GPX4 (Su et al. 2005) were described previously. All the other antibodies used in this study are commercially available as follows (ANT4, Abcam; P2X2, Santa Cruz; TOM20, Santa Cruz; TXNRD1, Bioss Antibodies; TXNRD2, Proteintech; acetylated tubulin, Sigma). All the chemicals were from Sigma Aldrich unless otherwise indicated.

### Single-Cell RNA-Seq Analysis

The raw count matrices for human (GSE109037) and mouse (GSE109033) testis single cell RNA (scRNA)-seq datasets (Hermann et al. 2018) were downloaded from Gene Expression Omnibus (GEO) database (https://www.ncbi.nlm.nih.gov/geo/). The downloaded raw count matrices were processed for quality control using the Seurat package (ver.3.2.3) (Stuart et al. 2019). Briefly, cells with less than 200 expressed features, higher than 9,000 (GSE109033) or 10,000 (GSE109037) expressed features and higher than 20% (GSE109037) or 25% (GSE109033) mitochondrial transcript fraction were excluded to select single cells with high quality mRNA profiles. The data was normalized by the total expression, scaled and log transformed. Identification of 2,000 highly variable features was followed by PCA to reduce the number of dimensions representing each cell. Statistically significant 15 PCs were selected based on the JackStraw and Elbow plots and provided as input for constructing a K-nearest-neighbors (KNN) graph based on the Euclidean distance in PCA space. Cells were clustered by the Louvain algorithm with a resolution parameter 0.1. Uniform Manifold Approximation and Projection (UMAP) was used to visualize and explore cluster data. Marker genes that define each cluster were identified by comparing each cluster to all other clusters using the MAST (Finak et al. 2015) provided in Seurat package. In order to correct batch effects among samples and experiments, we applied the Harmony package (ver.1.0) (Korsunsky et al. 2019) to the datasets. The Markov Affinity-based Graph Imputation of Cells (MAGIC) algorithm (ver.2.0.3) (van Dijk et al. 2018) was used to denoise and the count matrix and impute the missing data. In these testis datasets from adult human, we identified 23,896 high quality single cells, that were clustered into 7 major cell types, including spermatogonia, spermatocytes, early spermatids, late spermatids, peritubular myoid cell, endothelial cell, and macrophage. Similarly, we exploited 30,268 high quality single cells from eight adult and three 6-day postpartum mouse testis tissue samples and subsequently defined 11 major cell populations.

### Sperm Immunocytochemistry and Free Thiol Labeling Assay

Mouse and human sperm were washed in PBS twice, attached on the glass coverslips, and fixed with 4% paraformaldehyde (PFA) in PBS at room temperature (RT) for 10 min (mouse) or at 4°C for 1 hr (human). Fixed samples were permeabilized using 0.1% Triton X-100 in PBS at RT for 10 min, washed in PBS, and blocked with 10% goat serum in PBS at RT for 1 hr. Cells were stained with antibody in PBS supplemented with 10% goat serum at 4°C overnight. After washing in PBS, the samples were incubated with goat anti-rabbit Alexa 647 or Alexa 555-plus (Invitrogen, 1:1,000) in 10% goat serum in PBS at RT for 1 hr. Hoechst was used to counterstain nucleus for sperm head. For BODIPY-N-ethylmaleimide labeling, reduced thiols within proteins were alkylated with BODIPY FL maleimide (final concentration of 10 nM, ThermoFisher) for 30 min in the dark after permeabilization. The sample was then quenched by the addition of 500 mM 2-mercaptoethanol for 30 min in the dark, followed by 3 times washing in PBS. For ThiolTracker (ThermoFisher) labeling, fixed sperm were stained with 20 μM dye, followed by 3 times washing in PBS. Stained sperm samples were mounted with Prolong gold (Invitrogen) and cured for 24 hr, followed by imaging with Zeiss LSM710 Elyra P1 using Plan-Apochrombat 63X/1.40 or alpha Plan-APO 100X/1.46 oil objective lens (Carl Zeiss).

### Protein Extraction and Western Blotting

Whole sperm protein was extracted as previously described (Chung et al. 2011, Chung et al. 2014, Chung et al. 2017). In short, mouse epididymal spermatozoa washed in PBS were directly lysed in 2× SDS sample buffer or 8 M urea. The whole sperm lysate was centrifuged at 15,000 g, 4°C for 10 min. Supernatant were adjusted to 50 mM DTT and denatured at 95°C for 10 min before loading to gel. The membranes were incubated overnight at 4°C with primary antibody, followed by washing with PBST and incubation with appropriate secondary antibody for 1 hr. Bands were visualized using chemiluminescence and imaged by ChemiDoc (Bio-Rad). Antibodies used for Western blotting were rabbit anti-mouse TXNRD3 (1:2000), GPx4 (1:2000), ANT4, 1 μg/mL; P2X2, 1 μg/mL; TOM20, 0.2 μg/mL; TXNRD1, 1 μg/mL; TXNRD2, 1 μg/mL; acetylated tubulin and anti-acetylated tubulin (1: 20,000). Secondary antibodies were anti-rabbit IgG-HRP (1:10,000) and anti-mouse IgG-HRP (1:10,000) from Jackson ImmunoResearch (West Grove, PA).

### Protein Oxidation

Carbonyl groups inserted into proteins by oxidative reactions were evaluated with protein carbonyl assay kit (Abcam). Briefly, samples were homogenized and processed according to the manufacturer’s protocol. The carbonyl groups in the solubilized protein samples were derivatized using DNPH (2, 4 dinitrophenyl hydrazine) or control solution for 15 min and then neutralized. The samples were then loaded onto SDS PAGE gels and DNP conjugated proteins were detected by western blotting using primary DNP antibody and HRP conjugated secondary antibody.

### Flagella Waveform Analysis

To tether sperm head for planar beating, non-capacitated or capacitated spermatozoa (2-3×10^5^ cells) from adult male mice were transferred to the fibronectin-coated 37°C chamber for Delta T culture dish controller (Bioptechs) filled with HS and HEPES-buffered HTF medium (H-HTF)(Chung et al. 2017), respectively. Flagellar movements of the tethered sperm were recorded for 2 s with 200 fps using pco.edge sCMOS camera equipped in Axio observer Z1 microscope (Carl Zeiss). All movies were taken at 37°C within 10 min after transferring sperm to the imaging dish. FIJI software (Schindelin et al. 2012) was used to measure beating frequency of sperm tail, and to generate overlaid images to trace waveform of sperm flagella as previously described (Chung et al. 2017).

### Sperm Motility Analysis

Aliquots of sperm were placed in slide chamber (CellVision, 20 mm depth) and motility was examined on a 37°C stage of a Nikon E200 microscope under 10X phase contrast objective (CFI Plan Achro 10X/0.25 Ph1 BM, Nikon). Images were recorded (40 frames at 50 fps) using CMOS video camera (Basler acA1300-200um, Basler AG, Ahrensburg, Germany) and analyzed by computer-assisted sperm analysis (CASA, Sperm Class Analyzer version 6.3, Microptic, Barcelona, Spain). Sperm total motility and hyperactivated motility was quantified simultaneously. Over 200 motile sperm were analyzed for each trial, at least 3 biological replicates were performed for each genotype. To track swimming trajectory, the sperm motility was videotaped at 50 fps. The images were analyzed using Fiji software (Schindelin et al. 2012) by assembling overlays of the flagellar traces generated by hyperstacking binary images of 20 frames of 2 s movies coded in a gray intensity scale.

### Scanning Electron Microscopy

Sperm cells were attached on the glass coverslips and fixed with 2.5% glutaraldehyde (GA) in 0.1 M sodium cacodylate buffer (pH 7.4) for one hour at 4°C and post fixed in 2% osmium tetroxide in 0.1 M cacodylate buffer (pH 7.4). The fixed samples were washed with 0.1 M cacodylate buffer for 3 times and dehydrated through a series of ethanol to 100%. The samples were dried using a 300 critical point dryer with liquid carbon dioxide as transitional fluid. The coverslips with dried samples were glued to aluminum stubs and sputter coated with 5 nm platinum using a Cressington 208HR (Ted Pella) rotary sputter coater. Prepared samples were imaged with Hitachi SU-70 scanning electron microscope (Hitachi High-Technologies).

### Transmitted Electron Microscopy

Collected epididymal sperm cells were washed and pelleted by centrifugation and fixed in 2.5% GA and 2% PFA in 0.1 M cacodylate buffer pH 7.4 for one hour at RT. Fixed sperm pellets were rinsed with 0.1 M cacodylate buffer and spud down in 2% agar. The chilled blocks were trimmed, rinsed in the 0.1 M cacodylate buffer, and replaced with 0.1% tannic acid in the buffer for one hour. After rinsing in the buffer, the samples were post-fixed in 1% osmium tetroxide and 1.5% potassium ferrocyanide in 0.1 M cacodylate buffer for one hour. The post-fixed samples were rinsed in the cacodylate buffer and distilled water, followed by en bloc staining in 2% aqueous uranyl acetate for one hour. Prepared samples were rinsed and dehydrated in an ethanol series to 100%. Dehydrated samples were infiltrated with epoxy resin Embed 812 (Electron Microscopy Sciences), placed in silicone molds and baked for 24 hours at 60°C. The hardened blocks were sectioned in 60 nm thickness using Leica UltraCut UC7. The sections were collected on grids coated with formvar/carbon and contrast stained using 2% uranyl acetate and lead citrate. The grids were imaged using FEI TecnaiBiotwin Transmission Electron Microscope (FEI, Hillsbroro, OR) at 80 kV. Images were taken using MORADA CCD camera and iTEM (Olympus) software.

### Sperm Chromatin Dispersion Test (Halo Assay)

Samples were diluted to the concentration of 5-10×10^6^ cells/mL, then mixed with low-melting-point aqueous agarose to obtain a final concentration of 0.7% agarose. 50 μl mixture was dispensed onto a glass slide pre-coated with 0.65% standard agarose, then covered by a coverslip for solidification at 4°C for 4 min. The coverslip was then removed carefully before the sample was immersed into freshly prepared acid denaturation solution (0.08 N HCl) for 7 min, neutralizing and lysing solution 1 (0.4 M Tris-HCl, 0.8M DTT, 1% SDS, 50 mM EDTA, pH 7.5) for 10 min, and neutralizing and lysing solution 2 (0.4 M Tris-HCl, 2 M NaCl, 1% SDS, pH 7.5) for 5 min at RT in sequence. The slide was then transferred to Tris-borate EDTA solution (90 mM Tris-borate, 2 mM EDTA, pH 7.5) for 2 min, then dehydrated in sequential 70%, 90%, and 100% EtOH (2 min each). After drying, the slide was stained with DAPI (2 ug/ml), then examined under microscope as described above. The Halo size (diameter) was defined and classified into big (>17 μm), medium (between 12 and 17 μm) and small size (<12 μm).

### Acridine Orange Assay

Sperm cells were smeared on glass slide until dry, followed by washing in PBS pH 7.2. 4% paraformaldehyde (PFA) was used to fix cells for 15 min at RT, then covered by methanol for 5 min. washed with PBS for 5 min. The sample was incubated in RNAse A solution at 37°C for 30 min. Washing with PBS of pH 7.2 for 5 min was performed between each step. 0.1 M HCl was used for DNA denaturation for 30-45 sec, followed by AO staining working solution for 2 min. The slide was ready for examination under fluorescence microscopy after washing gently and drying.

### Mitochondrial Membrane Potential (ΔΨm) Measurement

Epididymal sperm were attached on Delta T culture chamber coated with poly-D-Lysine (2 mg/mL) or together with Cell-Tak for 30 min. Unattached sperm were removed by the gentle pipette wash. Mitotracker deep red (working concentration 500 nM, ThermoFisher) was loaded in sperm for 30 min, followed by gentle rinse with HS or HEPES-buffered HTF medium H-HTF. All movies were taken at 37°C by the pco.edge sCMOS camera equipped in Axio observer Z1 microscope (Carl Zeiss). The data were analyzed by Zen (Blue).

### Quantification and Statistical Analysis

Statistical analyses were performed using Student’s t-test or one-way analysis of variance (ANOVA) with Tukey post hoc test. Differences were considered significant at \**p* < 0.05, \*\**p* < 0.01, and \*\*\**p* < 0.001.

## Results

### *Txnrd3*^*-/-*^ Sperm Cells Display Defects in Morphology and Motility Under Capacitating Conditions

To understand why *Txnrd3*^*-/-*^ males exhibit sub-fertility *in vivo* and *in vitro* (Dou et al., 2021), we first examined sperm morphology and count from the cauda epididymis. The number of sperm produced by *Txnrd3*^*-/-*^ males was smaller than that from 2-3 months old heterozygous littermates (Supplementary Figure 1A). Among them, some *Txnrd3*^*-/-*^ sperm displayed hairpin formation within midpiece, specifically evident after 90 min incubation under capacitating conditions (Figure 1A, B), whereas sperm bent around annulus were negligible in both wild type and *Txnrd3*^*-/-*^ mice. Scanning electron microscopy (SEM) analysis did not reveal any gross abnormalities in the sperm midpiece (Figure 1C).

**Figure 1.**
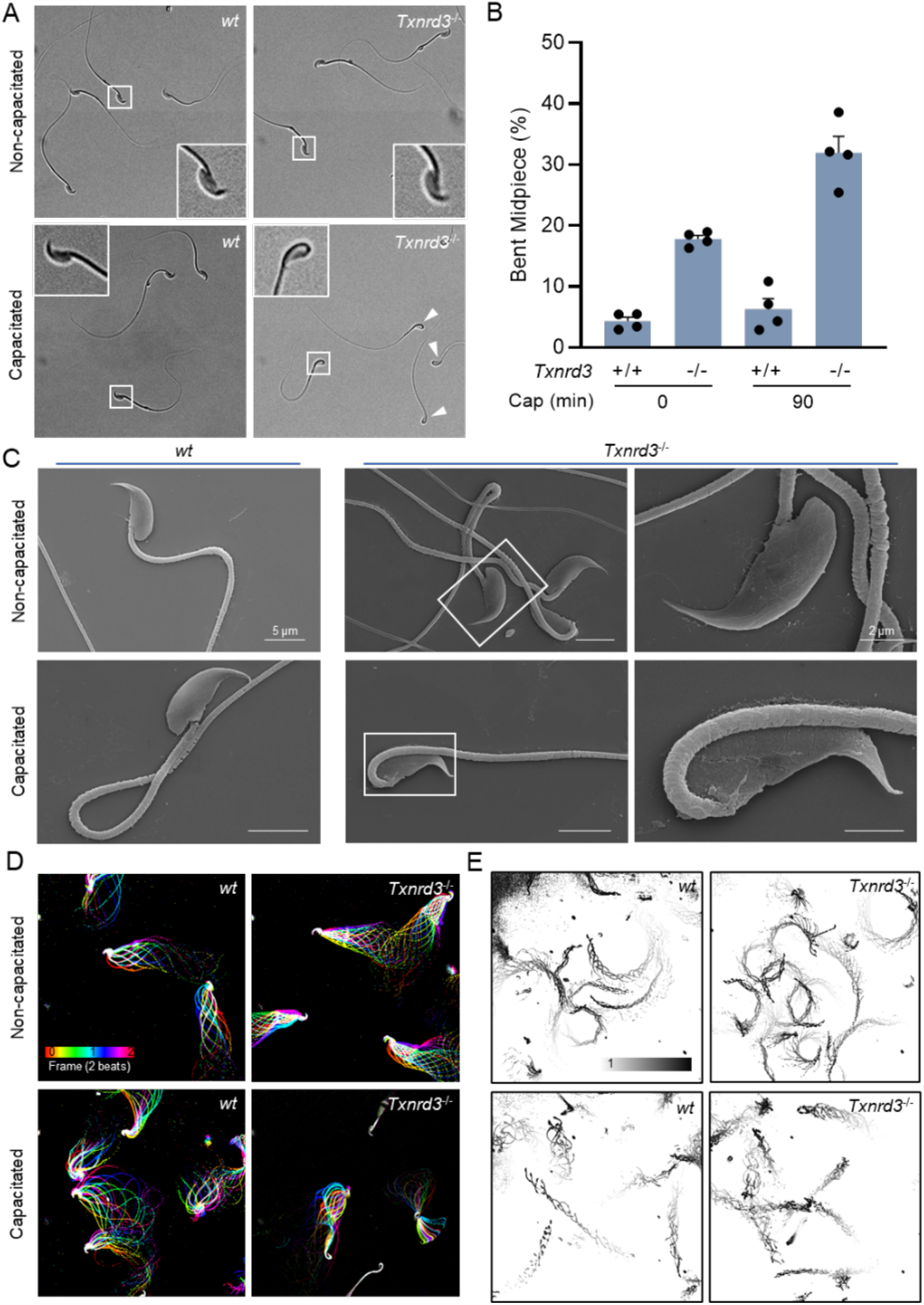
TXNRD3-deficient sperm are morphologically abnormal with impaired hyperactivated motility. (**A**) TXNRD3-deficient sperm exhibit abnormal hairpin morphology following 90 min incubation under capacitation conditions. (**B**) Quantification of abnormal hairpin morphology from **A**. Mean ± SEM, n = X each group. \*\**p* < 0.01 (**C**) Representative scanning electron microscopy (SEM) images of cauda epididymal sperm. (**D**) Aligned flagellar waveform traces. Movies recorded at 200 fps: *wt* and *Txnrd3*^-/-^ sperm cells attached on glass coverslips before (top) and after (bottom) 90 min capacitation. Overlays of flagellar traces from two beat cycles are generated by hyperstacking binary images; time coded in color. (**E**) Free swimming trajectory analysis. Head trace of free swimming *wt* and *Txnrd3*^-/-^ sperm cells at 0 min (top) and 90 min (bottom) after capacitation. Traces are from 1 s movies taken at 37°C.

Prompted by the observation of capacitation-associated hairpin formation within midpiece of *Txnrd3*^*-/-*^ sperm, we next analyzed sperm motility. Flagellar waveform analysis revealed that *Txnrd3*^*-/-*^ sperm become stiff in the midpiece and beat abnormally after incubating under capacitating conditions (Figure 1D, Supplementary Figure 1C and Video S1). Accordingly, CASA analysis found that overall motility of *Txnrd3*^*-/-*^ sperm was diminished (Supplementary Figure 1B). By contrast, hyperactivated motility was not significantly affected, likely because the *Txnrd3*^*-/-*^ sperm that are not bent still developed hyperactivated motility. To better understand the effect of TXNRD3 deficiency on free swimming pattern, sperm cells were placed in 0.3% methylcellulose which mimics viscous environment in the female reproductive tract. Swimming trajectory traced by time-lapse video microscope demonstrated that *Txnrd3*^*-/-*^ sperm move a shorter distance at a given time after capacitation (Figure 1E and Video S2), suggesting that the hairpin prevents them from swimming as efficiently as wild type sperm.

### Absence of TXNRD3 Affects the Redox State of Both Head and Tail of Sperm

TXNRD3 is enriched in the testis but not readily detected in the epididymis (Su et al. 2005). Our analysis of the publicly available scRNA-seq dataset (Speir et al. 2020) found that *Txrnd3* mRNA expression is high in spermatogonia and spermatocytes and reduced in spermatids (Supplementary Figure 2). We hypothesized that diminishing TXNRD3 mRNA and protein levels in epididymal sperm along the tract underlies the changes in the overall redox state of proteins during epididymal maturation and capacitation of sperm. Thus, we examined changes in TXNRD3 protein levels and found decreasing amount of TXNRD3 in sperm during their descent through epididymis (Figure 2A, B). We next examined the distribution and the extent of protein thiol modification in the absence of TXNRD3 by labeling epididymal sperm cells with ThiolTracker that reacts with reduced thiols (free “-SH”) irreversibly. Consistent with TXNRD3 levels from caput to cauda, wild type sperm from caput region exhibited strong fluorescence over the entire cell whereas corpus and cauda sperm displayed gradually diminishing fluorescence intensity, more noticeably in the head (Figure 2C, D). These results are in line with the previously reported increasing disulfide bond formation in sperm during their transit along the male reproductive tract (Dias et al. 2014).

**Figure 2.**
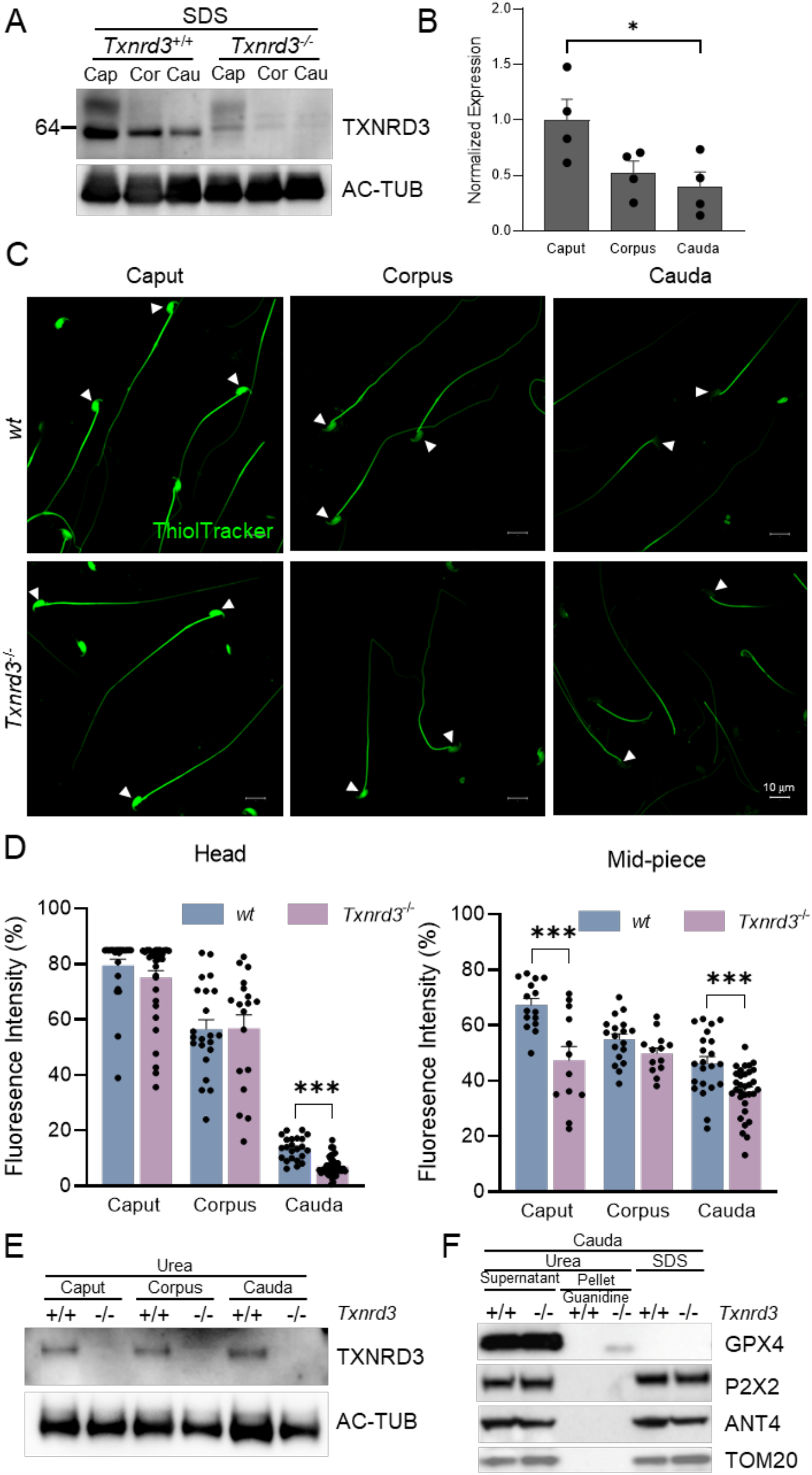
Free thiol level and oxidative status are altered in *Txnrd3*^-/-^ spermatozoa. (**A**) TXNRD3 expression decreased in concert with sperm maturation in the epididymis. (**B**) Quantification of TXNRD3 expression from (A). (**C**) Free thiol groups level in epididymal sperm from *wt* and *Txnrd3*^-/-^ mice. Spermatozoa isolated from the caput, corpus and cauda epididymis were incubated with ThiolTracker. (**D**) Relative quantification of free thiol levels in sperm head and midpiece. (**E**) TXNRD3 are equally extracted by urea from cauda sperm. (**F**) GPx4, a potential TXNRD3 substrate, exhibited stronger resistance to lysis buffer in the absence of TXNRD3. Other mitochondrial or midpiece localized proteins such as ANT4, P2X2 and TOM20 were well solubilized in all detergents tested.

Using ThiolTracker or BODIPY-NEM, another thiol-reactive dye, we found that, in the absence of TXNRD3, free thiol content is further decreased modestly, yet significantly, in cauda sperm (Figure 2C, D and Supplementary Figure 3A). It is possible that other reductases function redundantly and compensate for the loss of TXNRD3; e.g. both TXNRD1 and TXNRD2 proteins are detected in wild type and *Txnrd3*^*-/-*^ sperm (Supplementary Figure 3D), and there are other thiol oxidoreductases, including those specific to the male reproductive system, that are also expressed. As sperm proteins were found more oxidized in cauda than in caput, we speculate that the efficiency of protein extraction can be affected by the overall redox status. Indeed, when we used urea instead of SDS to examine protein levels of TXNRD3 from caput, corpus, and cauda sperm, we did not observe significant difference (Figure 2E), suggesting that TXNRD3 itself is likely a substrate of thioredoxin system during epididymal maturation.

Sperm midpiece harbors compartmentalized mitochondria (Skinner 2018) (see also Figure 1C). As *Txnrd3*^*-/-*^ sperm specifically exhibit the hairpin formation within the midpiece, next we tested whether the TXNRD3 loss-of-function affects the redox states of mitochondrial proteins more specifically. In the SDS extracted fraction, the protein level of glutathione peroxidase 4 (GPX4) -a mitochondrial structural protein and potential substrate of TXNRD3 (Ursini et al. 1999, Su et al. 2005) – was much lower in the cauda sperm just as seen for TXNRD3 (Figure 2A, B and Supplementary Figure 3B,C). By contrast, urea readily extracted GPx4 even from cauda sperm (Figure 2F, *supernatant*), indicating that GPx4 becomes likely more oxidized and therefore SDS-insoluble during epididymal transit. Intriguingly, we observed a urea-insoluble GPx4 fraction not in wild type but in *Txnrd3*^*-/-*^ cauda sperm when the pellet was further solubilized by guanidine following urea treatment (Figure 2F, *pellet*). Other non-structural but membrane proteins localized in sperm mitochondria or midpiece such as ANT4, P2X2 and TOM20 and the other two thioredoxin reductases -TXNRD1 and TXNRD2 -were well solubilized in all detergents tested (Figure 2F and Supplementary Figure 3D). Taken together, our data clearly showed that TXNRD3 is involved in redox regulation of sperm proteins during epididymal transition.

### Capacitation-Associated Oxidation is Deregulated in *Txnrd3*^*-/-*^ Sperm (Chromatin Organization)

It has been known that redox signaling is involved in sperm capacitation (O’Flaherty 2015). We next examined how free thiol levels change during sperm capacitation. Under capacitating conditions, wild type sperm were less reactive with ThiolTracker, particularly in the head, whereas *Txnrd3*^*-/-*^ sperm was not reactive regardless (Figure 3A). These results suggest that sperm proteins were more oxidized in the absence of TXNRD3 during sperm maturation, which remain relatively stable during capacitation. To test this idea, we examined protein oxidation levels by detecting carbonyl groups inserted into proteins. Incubating sperm under capacitating conditions dramatically increased sperm protein carbonylation, suggesting active oxidative reactions occur. The protein oxidation level was further enhanced in *Txnrd3*^*-/-*^ sperm (Figure 3B and Supplementary Figure 4).

**Figure 3.**
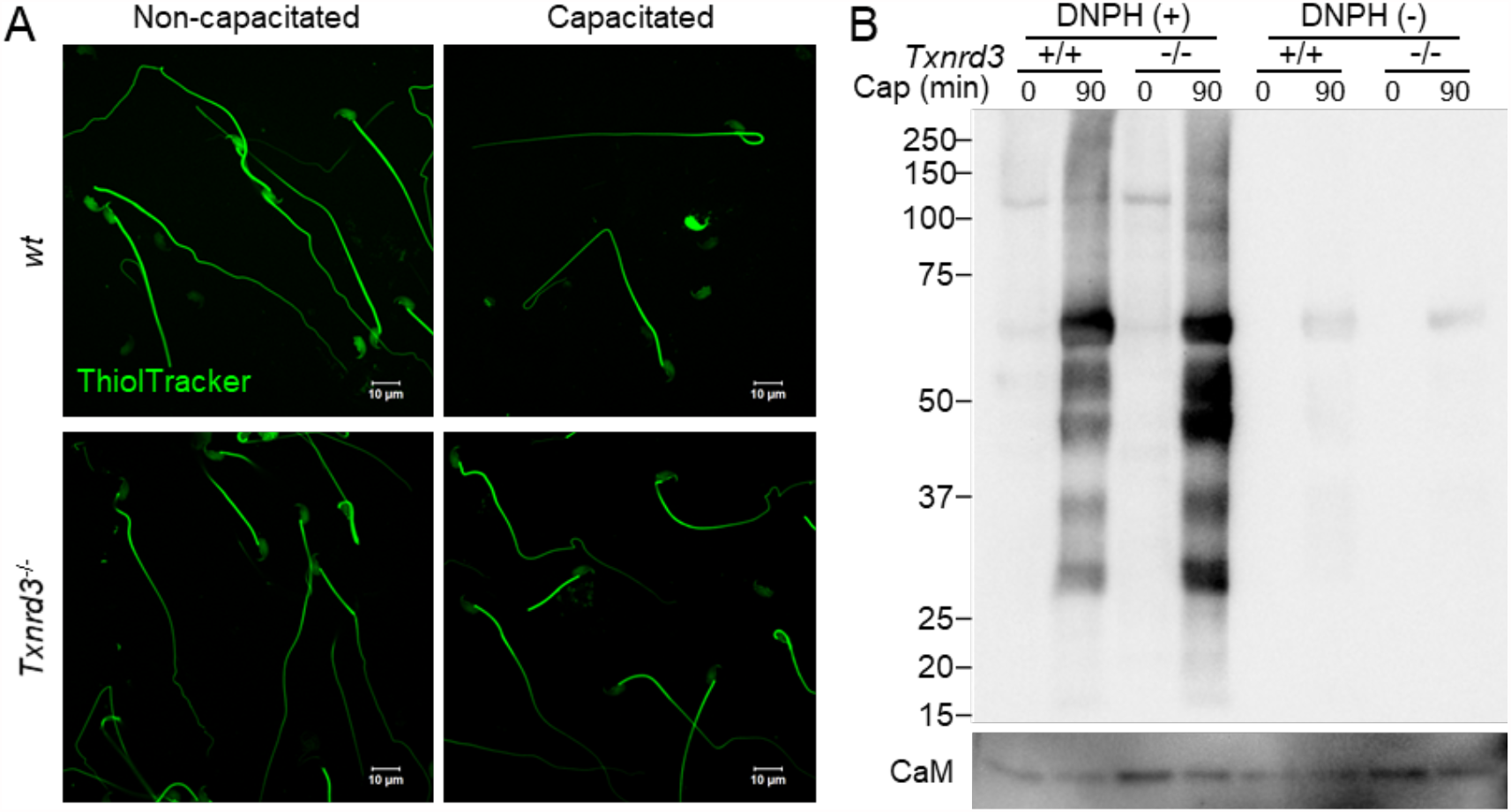
TXNRD3-deficient sperm undergo more oxidation after capacitation. (**A**) Free thiol groups level revealed with ThiolTracker in cauda epididymal sperm from wide type and *Txnrd3*^-/-^ mice before and after 90 min capacitation. (**B**) Protein carbonylation level in *wt* and *Txnrd3*^-/-^ sperm before and after capacitation. The carbonyl groups derivatized by DNPH (2, 4 dinitrophenyl hydrazine) were detected by western blotting using DNP antibody.

As the free thiol level changes were most dramatic in the head, we next investigated the loss of TXNRD3 function on sperm chromatin organization. We examined the extent of DNA integrity by halo assay where radial dispersion of the DNA fragments from the nucleus was measured upon artificial acid denaturation; the bigger the halo size is, the less damaged DNA fragment is present (Fernández et al. 2003, Chohan et al. 2006, Tandara et al. 2014). Intriguingly, *Txnrd3*^*-/-*^ sperm resist the denaturation better, developing smaller halos than those of wild type sperm (Figure 4). This result suggests that more damaged, double stranded DNA fragments are present in the nucleus in the absence of TXNRD3, likely due to more DNA fragments crosslinked to proteins. The nuclei of sperm incubated under capacitating conditions, developed smaller halos in both wild type and *Txnrd3*^*-/-*^ sperm (Figure 4). We further evaluated the altered resistance of DNA to acid denaturation by acridine orange (AO) assay; AO produces different emission when bound to single vs. double stranded DNA. We observed that double-stranded DNA was more stable and resistant to acid denaturation in both nucleus and mitochondria in capacitated *Txnrd3*^*-/-*^ sperm (Supplementary Figure 5). Taken together, the thiol oxidoreductase function is diminished in *Txnrd3*^*-/-*^ sperm compared to wild type sperm. The data show that TXNRD3 supports redox regulation of epididymal sperm maturation, protecting mitochondrial structure and chromatin from oxidative damage during sperm capacitation.

**Figure 4.**
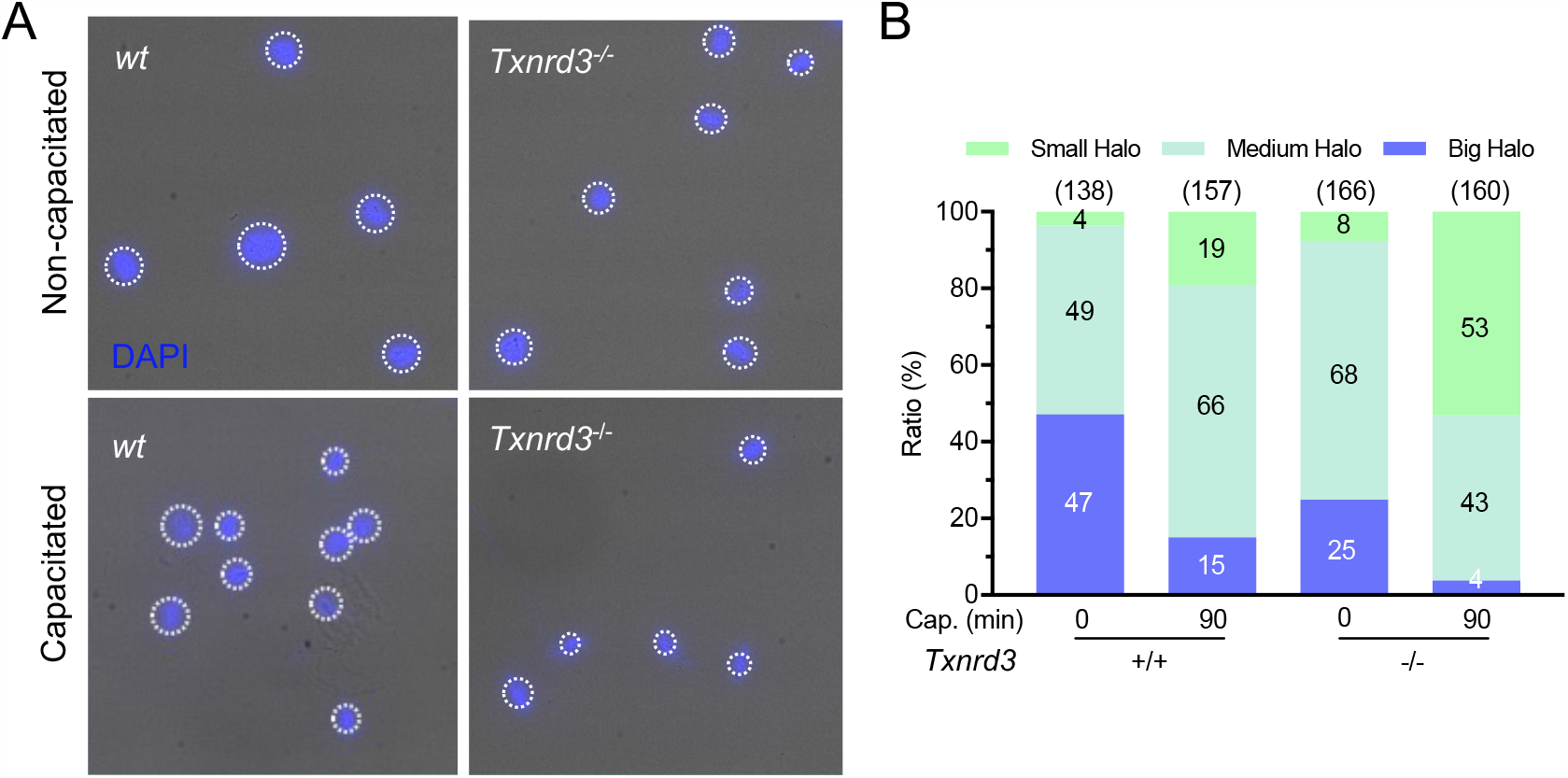
TXNRD3-deficient sperm are more stable and resistant to acid denaturation after capacitation. (**A**) Sperm chromatin dispersion test show smaller halo in *Txnrd3*^-/-^ sperm than in *wt* sperm during capacitation, indicating extensive fragmented DNA after acid denaturation during the assay procedure. DAPI (Blue). (**B**) Quantification of various halo size from *wt* and *Txnrd3*^-/-^ sperm before and after capacitation. Based on sperm head size (∼8 μm), halo sizes which are <12 μm (50% larger), <17 μm (100% larger) and >17 μm are divided into small, medium and big, respectively.

### Lack of TXNRD3 Dysregulates Capacitation-Induced Remodeling of Sperm Mitochondrial Ultrastructure

To maintain motility and support capacitation after ejaculation, mammalian sperm utilize nutrient molecules in the seminal plasma and in the female reproductive tract environment to generate energy (Visconti 2012). As terminally differentiated and highly polarized cells, sperm uniquely compartmentalize metabolic pathways in the flagella: the glycolytic enzymes are specifically localized in the principal piece, whereas mitochondrial oxidative phosphorylation (OXPHOS) occurs in the midpiece (Skinner 2018). A classic view on mouse sperm bioenergetics has been that glycolysis is the main metabolic pathway to support tyrosine phosphorylation and hyperactivation whereas both glycolytic and OXPHOS are functional (Travis et al. 2001, Mukai and Okuno 2004, Goodson et al. 2012). Intriguingly, our transmission electron microscopy (TEM) analysis revealed that mitochondria display more well-defined cristae after incubating under capacitating conditions (Figure 5A, B). Dimerization of the V-like shaped mitochondrial ATPase and its self-assembly into rows drives the cristae organization and determines respiratory efficiency (Dudkina et al. 2006, Strauss et al. 2008, Cogliati et al. 2013, Blum et al. 2019). Thus, our results suggest that capacitation increases mitochondrial activity in mouse sperm. In addition, we observed that capacitation induces electron-dense cores in the mitochondrial matrix of capacitated sperm cells (Figure 5A, B). Notably, *Txnrd3*^*-/-*^ sperm developed significantly more dense cores than wild type sperm regardless of capacitation status. As similar dense cores were previously reported in *Pmca4*^-/-^ sperm that might potentially experience Ca^2+^ overload (Okunade et al. 2004), this result suggests that the TXNRD3-dependent redox control might impact on Ca^2+^ signaling during sperm capacitation.

**Figure 5.**
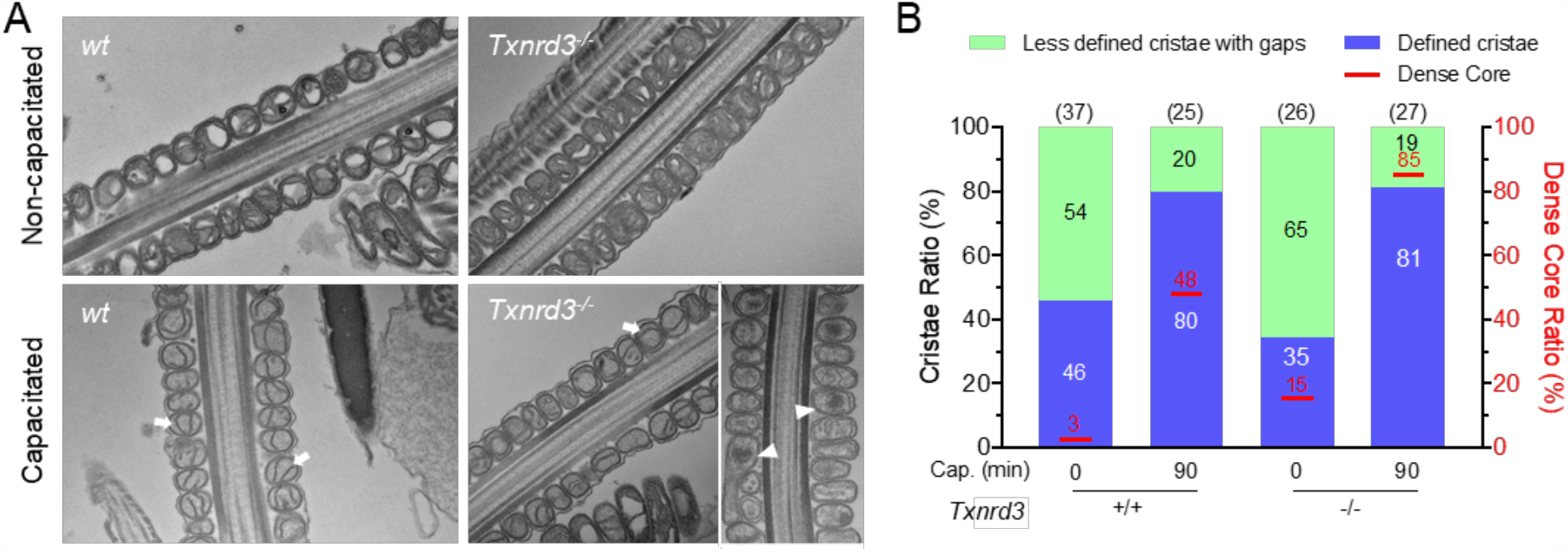
Ultrastructural analysis of epididymal sperm. (**A**) Representative transmission electron microscopy (TEM) images of cauda epididymal sperm mitochondria. Arrows indicate defined cristae found in mitochondria of capacitated sperm; arrow heads indicate dense cores found in mitochondria of *Txnrd3*^-/-^ sperm. (**B**) Classification of mitochondrial morphology in *wt* and *Txnrd3*^-/-^ sperm before and after capacitation.

### *Txnrd3*^*-/-*^ Sperm Lose Control of Mitochondrial Membrane Hyperpolarization

Mitochondrial membrane potential (ΔΨm) is an essential driving force for oxidative phosphorylation. To better understand how mitochondrial activity changes during sperm capacitation and the loss-of-function effect of TXNRD3 on this process, we performed functional imaging to probe ΔΨm using MitoTracker DeepRed, a fluorescent dye specifically taken up by mitochondria in live cells according to ΔΨm (Lugli et al. 2005, Zhou et al. 2011, Greene et al. 2012). The decreased fluorescence intensity of MitoTracker DeepRed indicates mitochondrial depolarization. In agreement with the higher mitochondrial activity during capacitation, dissipated ΔΨm by antimycin A -an inhibitor of oxidative phosphorylation -was greater in wild type sperm incubated under capacitating conditions (Figure 6 and Supplementary Figure 6). Under non-capacitating conditions, we find that ΔΨm is not significantly different between wild type and *Txnrd3*^*-/-*^ sperm. Under capacitating conditions, however, we found two distinct populations among TXNRD3-deficient sperm in terms of ΔΨm (Figure 6 and Supplementary Figure 6); one group did not show any change in the fluorescent intensity of the dye upon antimycin A, suggesting reduced or no mitochondrial activity (red trace) whereas the other group exhibited ΔΨm at a similar level to that of wild type sperm. These results further show that the loss of TXNRD3 dysregulates the capacitation-associated remodeling of mitochondrial function in mouse sperm.

**Figure 6.**
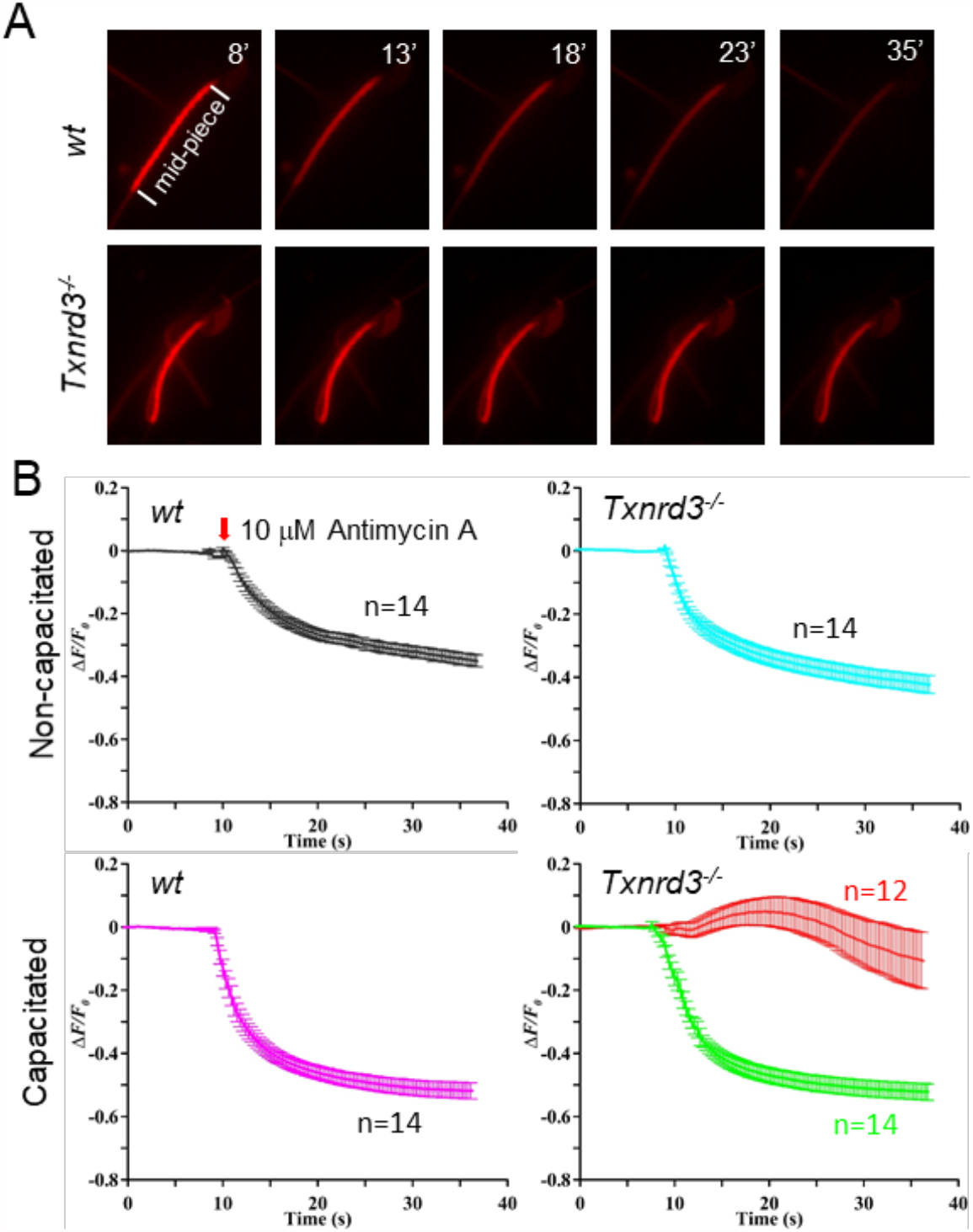
Mitochondrial activity, indicated by membrane potential, is impaired in absence of TXNRD3. (**A**) Dynamic representative of mitochondrial membrane potential (ΔΨm) indicated by Mitotracker. Mouse sperm were stained with 500 nM Mitotracker Deep Red which accumulates ΔΨm-dependently. (**B**) Transition of the mitochondrial membrane potential. The changes of fluorescence intensity were calculated as ΔF/F_0_ (F_0_, the mean fluorescence intensity of the sperm midpiece before adding Antimycin A (at 10 s), an inhibitor of electron transport chain; F, the fluorescence of the midpiece after adding antimycin A; ΔF=F-F_0_). Capacitated *Txnrd3*^-/-^ sperm show heterogeneous response to antimycin A, suggesting disrupted cellular respiration.

## Discussion

### Evidence that TXNRD3 Supports Sperm Function and Redox Regulation in Male Fertility

In the accompanying manuscript, we show that *Txnrd3*^*-/-*^ male mice exhibit reduced fertility *in vivo* and *in vitro* (Dou et al., 2021). The findings in this study demonstrate that the subfertility likely results from the combinatorial effects of defective chromatin organization and capacitation-induced tail bending within the midpiece. The high degree of sperm DNA fragmentation was reported to correlate with low fertilization rates, embryo quality and pregnancy after IVF and intracytoplasmic sperm injection (Loft et al. 2003, Tandara et al. 2014). The bent tail would force the head backwards, opposite to the swimming direction of the sperm. Additionally, TXNRD3 in sperm is gradually oxidized during epididymal transit, which is consistent with reduced free thiol group levels in cells along the epididymal tract. Therefore, the main timing of TXNRD3 function is during spermatogenesis in the testis and epididymal maturation rather than the subsequent fertilization process. Moderate changes in the free thiol levels and redox status of the *Txnrd3*^*-/-*^ sperm may be due to functional redundancy of other thiol reductases that partly compensate for the loss of TXNRD3. The data are also consistent with the mode wherein TXNRD3 supports accelerated redox remodeling during sperm maturation, which may also occur, albeit at a lower rate, in the absence of this enzyme. Alternatively, the overall redox status is only marginally changed in the absence of TXNRD3, but the pattern of disulfide bond formation could be jumbled, sufficient to make *Txnrd3*^*-/-*^ sperm vulnerable to capacitation-associated oxidative damage.

### Mitochondrial Function and Control of Metabolism During Capacitation

Mitochondria is a key organelle still preserved in the fully differentiated spermatozoa. Previous knock-out studies of sperm-specific glyceraldehyde 3-phosphate dehydrogenase and phosphoglycerate kinase 2 have identified glycolysis as the main source of ATP to sustain the motility of mouse spermatozoa (Miki et al. 2004, Danshina et al. 2010). Because OXPHOS yields more ATP per mol of glucose than glycolysis, the discordant contribution of the two metabolic pathways to provide the energy for mouse sperm motility has been an unresolved puzzle for a long time. A comparative study revealed that sperm of closely related mouse species with higher oxygen consumption rate were able to produce higher amounts of ATP, achieving higher swimming velocities (Tourmente et al. 2015).

Here we report, to our knowledge for the first time, that capacitation induces sperm mitochondria to form more defined cristae (Figure 5). As ATPase dimerization and self-assembly drive cristae formation (Dudkina et al. 2006, Strauss et al. 2008, Blum et al. 2019), this result indicates that capacitation enhances OXPHOS pathway to supply additional ATP. The higher metabolic activity in capacitating sperm is further supported by our observation of a more hyperpolarized mitochondrial membrane potential (ΔΨm) (Figure 6), an indication of protonmotive force and mitochondrial function. Consistently, a recent study reported accelerated oxygen consumption rate in capacitated sperm (Balbach et al. 2020).

Interestingly, we observed that a significant portion of TXNRD3-deficient sperm do not only fail to maintain ΔΨm but also develop a more apparent electron-dense core in the mitochondrial matrix. As a condensed mitochondrial matrix was observed in the absence of plasma membrane Ca^2+^-ATPase 4 (Okunade et al. 2004), this result suggests that calcium is overloaded in *Txnrd3*^*-/-*^ sperm mitochondria during sperm capacitation. Whether the TXNRD3-deficient sperm cells that cannot maintain ΔΨm are the same sperm populations with the electron-dense core requires further work. The small cytosol volume in mature sperm makes it difficult to achieve spatial resolution specific to mitochondria Ca^2+^ imaging with the currently available Ca^2+^ dyes. Generation of a mouse model expressing a genetically encoded Ca^2+^ indicator specifically targeted to sperm mitochondria will help to directly address the functional relationship of redox regulation of mitochondria dynamics in Ca^2+^ homeostasis and sperm metabolism during capacitation.

In summary, our study investigated the TXNRD3 function and potential cellular mechanisms in sperm maturation and male fertility. The absence of TXNRD3 deregulates the redox control of sperm chromatin and mitochondrial structure proteins during capacitation, resulting in distinct hairpin-like bending and defective mitochondrial respiration; the loss-of-function prevents *Txnrd3*^*/-*^ sperm from properly positioning to penetrate the egg, compromising efficient ATP production. *Txnrd3*^*-/-*^ mice are fertile in controlled laboratory environment without limitation of resource and male competition. Yet our findings from *in vitro* experiments show that TXNRD3 and the thioredoxin system intimately contribute to sperm function and male reproduction. This new knowledge could help to define conditions for assisted reproductive technology in clinic.

## Conflict of Interest

The authors declare no competing interests.

## Author Contributions

J.-J.C. and V.N.G. conceived the study. J.-J.C. oversaw the project. J.-J.C. and H.W. designed the experiments. H.W., Q.D., and J.-J.C. performed experiments. K.J. and J.C. analyzed scRNAseq database. H.W. prepared the figures and H.W. and J.-J.C wrote the initial draft of the manuscript. H.W., V.N.G. and J.-J.C. edited the manuscript with input from all other authors in the final version.

## Funding

This work was funded by NIH R01GM065204 to V.N.G, and the start-up funds from Yale School of Medicine and NIH R01HD096745 to J.-J.C.

## Acknowledgments

We thank the Harvard Medical School EM facility (TEM) and the Yale Center for Cellular and Molecular Imaging (SEM) for assistance in electron microscopy, Jae Yeon Hwang for help in analyzing TEM images of sperm mitochondria, and Xiaofang Huang for initial analysis of free thiol measurement.

---

**Supplementary Figure 1.**
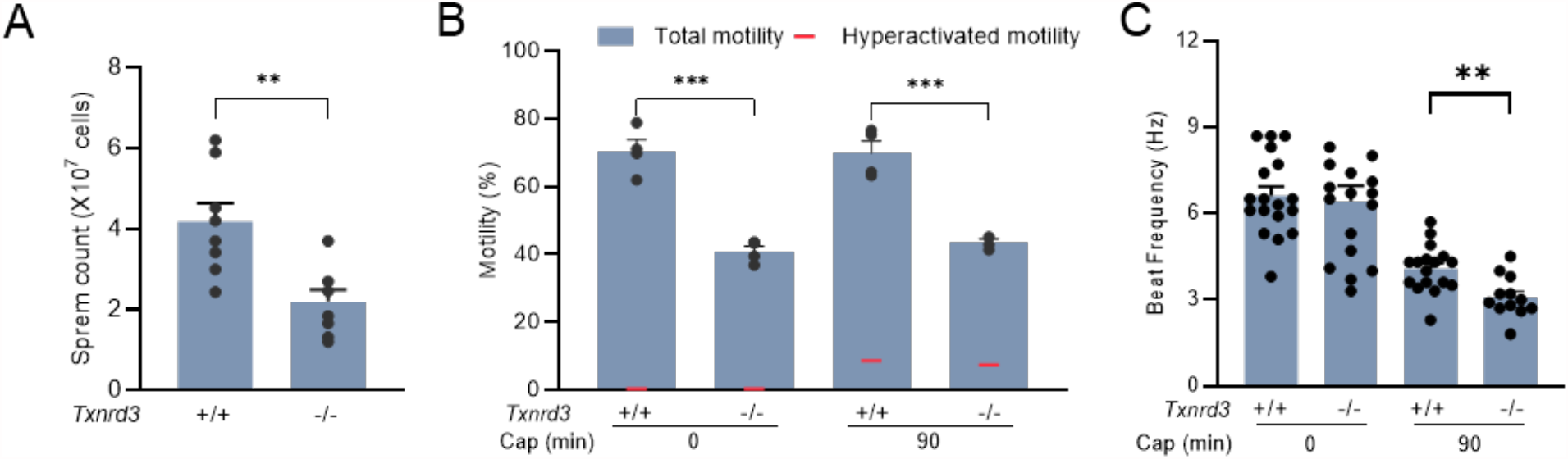
TXNRD3-deficient mice exhibit reduced sperm count and motility. (**A**) Epididymal sperm count (Mean ± SEM) from littermates at ages of 3-4 months of age. (**B**) Sperm total motility and hyperactivated motility before and after incubating under capacitation conditions. (**C**) Flagellar beat frequency of *wt* and *Txnrd3*^-/-^ sperm before (0 min, *wt*, 6.6 ± 0.3 Hz; *Txnrd3*^-/-^, 6.4 ± 0.5 Hz) and after (90 min, *wt*, 4.1 ± 0.2 Hz; *Txnrd3*^-/-^, 3.1 ± 0.2 Hz) capacitation. Mean ± SEM, n = X each group. \*\**p* < 0.01, \*\*\**p* < 0.001.

Supplementary Video 1. Head-tethered sperm from *wt* and *Txnrd3*^-/-^ mice before and after incubating under capacitation conditions.

upplementary Video 2. Free-swimming sperm from *wt* and *Txnrd3*^/-^ mice before and after incubating under capacitation conditions.

**Supplementary Figure 2.**
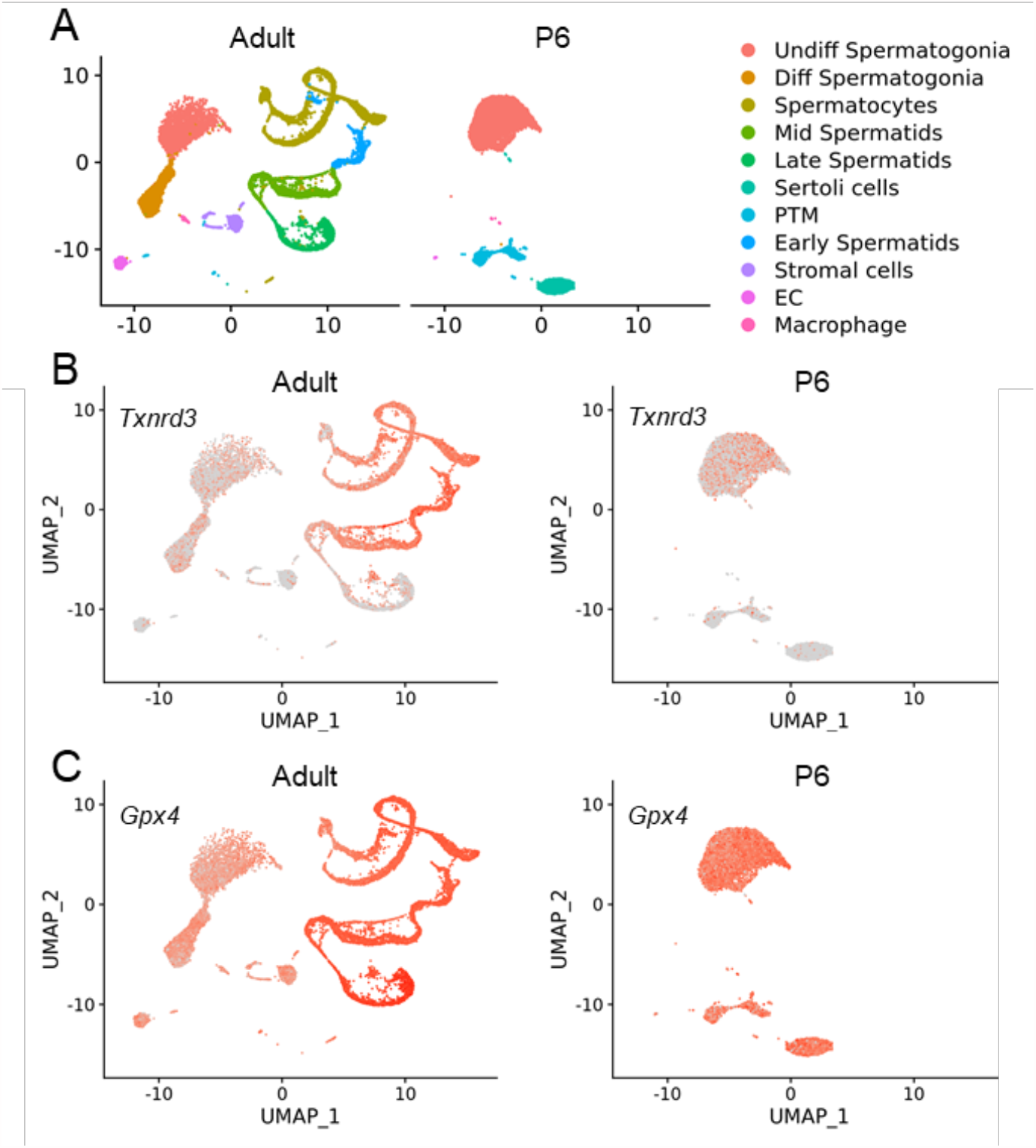
Single testicular cell transcriptome profiling from adult and postnatal day 6 (P6) mice. (**A**) Clustering analysis of combined single-cell transcriptome data from testes. Each dot represents a single cell and is colored according to its cluster identity as indicated on the figure key. 11 cluster identities were assigned based on marker gene expression. PTM, Peritubular myoid; EC, Testicular endothelial cells. (**B**) Expression patterns of *Txnrd3* which is highly expressed in spermatogonia and spermatocytes but much less in spermatids. Red indicates high expression and gray indicates low or no expression. (**C**) Expression patterns of *Gpx4*, which is abundant through spermatogenesis.

**Supplementary Figure 3.**
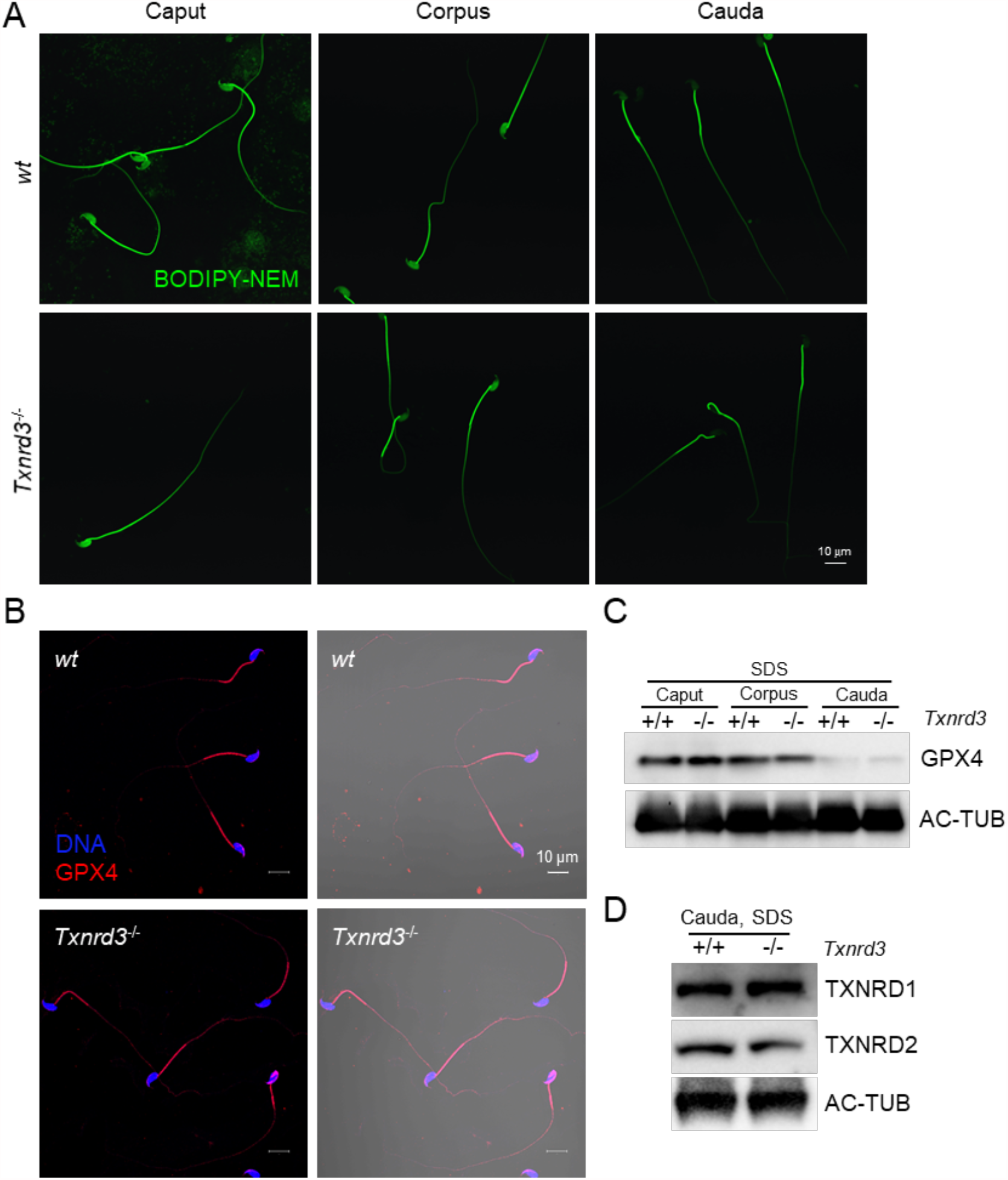
Free thiol level and level of GPX4 from caput to cauda sperm. (**A**) Spermatozoa isolated from the caput, corpus and cauda epididymis were incubated with Bodipy-labeled NEM. After quenching with β-mercaptoethanol, free thiol groups in the cells were visualized using confocal microscope. (**B**) Localization of GPx4 (**C**) GPx4 shows descending level in sperm during epididymal transit. (**D**) TXNRD1 and TXNRD2 show same level in both wide type and *Txnrd3*^-/-^ sperm.

**Supplementary Figure 4.**
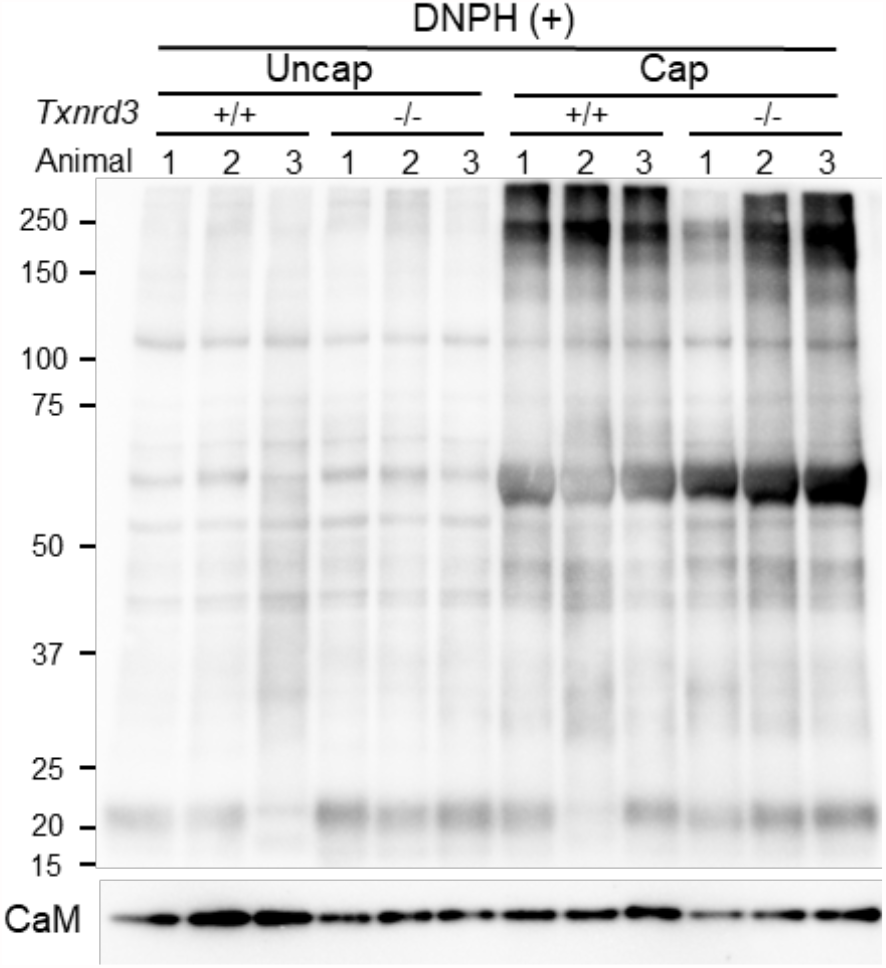
Protein oxidation status probed by protein carbonylation level is changed during sperm capacitation. Protein carbonylation level was detected in *wt* and Txnrd3-/-sperm before and after capacitation from multiple mice.

**Supplementary Figure 5.**
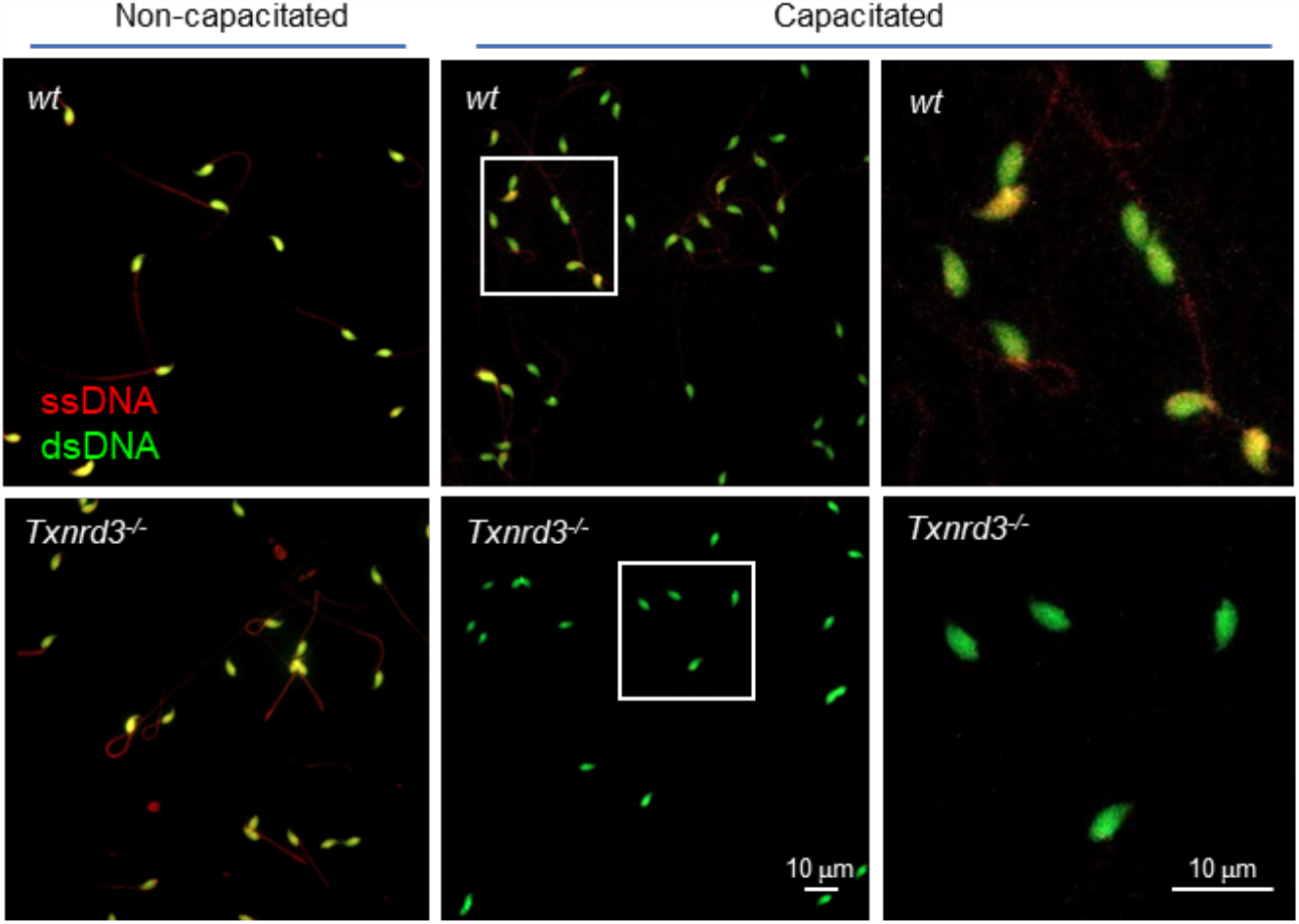
Evaluation of DNA state by Acridine Orange assay. After HCl incubation, *Txnrd3*^-/-^ sperm generated less single-stranded DNA after capacitation, indicating more resistance to acid denaturation. Single-stranded DNA (Red), double-stranded DNA (Green).

**Supplementary Figure 6.**
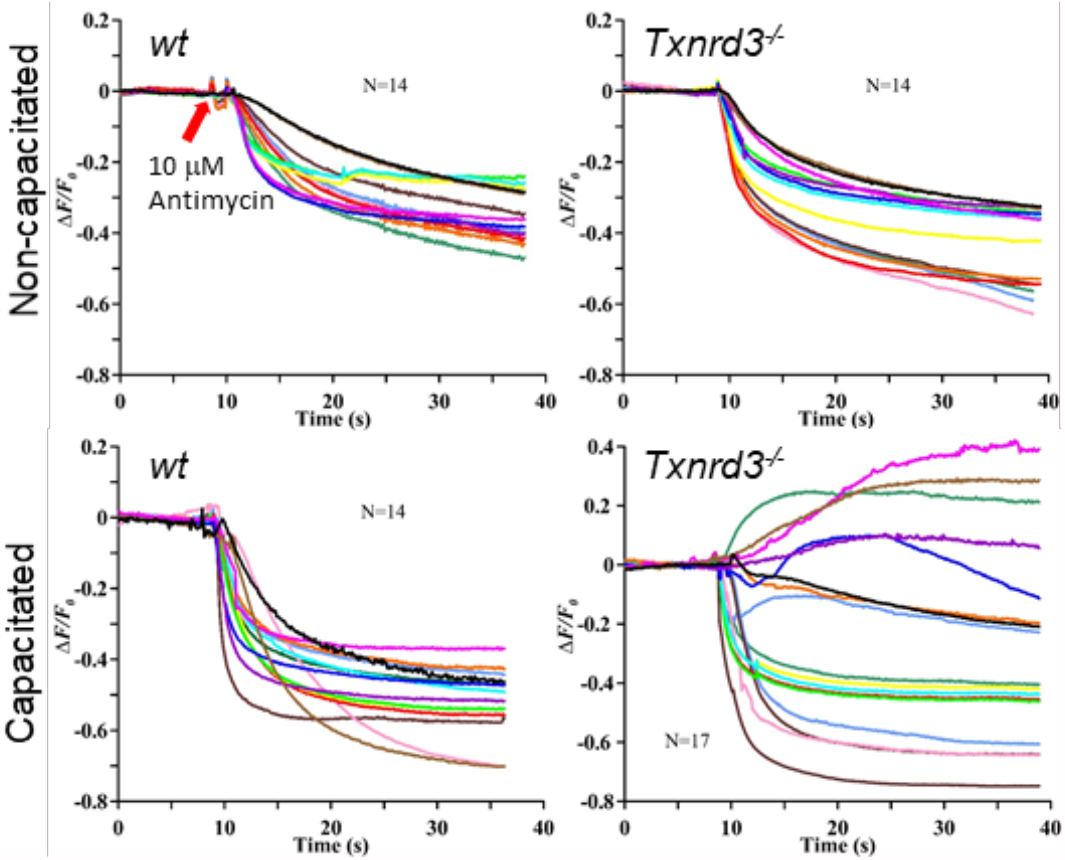
Individual traces of mitochondrial membrane potential transition from wild type and *Txnrd3*^-/-^ sperm. Presented are the traces of the individual sperm cells for Figure 6B.

## Notes

### Competing Interest Statement

The authors have declared no competing interest.

